# CoVaccine HT™ adjuvant potentiates robust immune responses to recombinant SARS-CoV-2 Spike S1 immunisation

**DOI:** 10.1101/2020.07.24.220715

**Authors:** Brien K. Haun, Chih-Yun Lai, Caitlin A. Williams, Teri Ann Wong, Michael M. Lieberman, Laurent Pessaint, Hanne Andersen-Elyard, Axel T. Lehrer

**Affiliations:** Department of Tropical Medicine, John A. Burns School of Medicine, University of Hawaii, Honolulu, HI; Cell and Molecular Biology Graduate Program, John A. Burns School of Medicine, University of Hawaii, Honolulu, HI; BIOQUAL, Inc., Rockville, MD.

**Author notes:** **Corresponding author**: Axel T. Lehrer, Dr. rer. nat., Assistant Professor, Department of Tropical Medicine, John A. Burns School of Medicine, University of Hawaii, Honolulu, HI, 651 Ilalo St., Biomedical Sciences Building, Honolulu HI 96813.

**Keywords:** SARS-CoV-2, COVID-19, Adjuvant, Vaccines, Preclinical vaccine studies.

## Abstract

The current COVID-19 pandemic has claimed hundreds of thousands of lives and its causative agent, SARS-CoV-2, has infected millions, globally. The highly contagious nature of this respiratory virus has spurred massive global efforts to develop vaccines at record speeds. In addition to enhanced immunogen delivery, adjuvants may greatly impact protective efficacy of a SARS-CoV-2 vaccine. To investigate adjuvant suitability, we formulated protein subunit vaccines consisting of the recombinant S1 domain of SARS-CoV-2 Spike protein alone or in combination with either CoVaccine HT™ or Alhydrogel. CoVaccine HT™ induced high titres of antigen-binding IgG after a single dose, facilitated affinity maturation and class switching to a greater extent than Alhydrogel and elicited potent cell-mediated immunity as well as virus neutralising antibody titres. Data presented here suggests that adjuvantation with CoVaccine HT™ can rapidly induce a comprehensive and protective immune response to SARS-CoV-2.

## 1. Introduction

The outbreak of 2019-novel coronavirus (SARS-CoV-2), the etiological agent of Coronavirus Disease 2019 (COVID-19), began in Wuhan, China in late 2019 and quickly spread across the globe causing epidemics on every continent except Antarctica in under four months. This virus has caused more than 15.5 million cases and over 630,000 deaths worldwide (as of 7-24-20) ^1^. SARS-CoV-2 is highly transmissible during both the pre-symptomatic and acute symptomatic phases and the infection fatality rate has been reported as high as 3.4% ^2^. COVID-19 often develops into severe illness, including pneumonia. Currently, there are no licensed vaccines or effective therapeutic strategies available to treat COVID-19. Evidence indicates that high titres of antibody targeting the Spike protein may neutralise virus, a concept which carries credence for the closely related SARS-CoV ^3–5^. Vaccine platforms at the forefront of development are mRNA-based, DNA-based, virally vectored (replication competent or incompetent), as well as recombinant protein subunits ^6,7^. Many vaccine candidates may require adjuvantation to induce robust immune responses and rapidly induce high antibody titres. However, no consensus is established on an optimal adjuvant that best induces protective immunity to SARS-CoV-2.

In order to investigate which adjuvants induce a strong humoral response, our group has formulated protein subunit vaccine candidates using a recombinant SARS-CoV-2 Spike subdomain 1 (S1) protein, obtained from Sino Biological, Inc., adjuvanted with either CoVaccine HT™ or Alhydrogel. The former is a proprietary adjuvant and the latter is an FDA approved adjuvant used in several FDA licensed vaccines. CoVaccine HT™ is an oil-in-water emulsion of hydrophobic, negatively-charged sucrose fatty acid sulphate esters with the addition of squalane^8,9^ whereas Alhydrogel is a colloid of aluminium hydroxide which binds protein to facilitate antigen recognition and thus, improve the immune response ^10^. The mechanism of action of Alhydrogel remains somewhat elusive, however this adjuvant likely interacts with NOD like receptor protein 3 (NLRP3) but does not interact with TLRs^11^. This difference in cellular activation can account for the disparities seen between the use of CoVaccine HT™ and Alhydrogel presented here. The stabilised oil in water emulsion functions by generating a response skewed towards a T-helper type 1 cell (T_h_1) direction which can in turn sustain CD8 T cells capable of mitigating viral infection^12^. This adjuvant is also capable of inducing T cell differentiation to T-follicular helper (T_fh_) cells which is evident through immunoglobulin class switching to IgG2a^13^. In concert, these cellular responses enhance the humoral response evidenced by the overall higher titres of IgG^13^.

CoVaccine HT™ also offers an advantage in comparison to Alhydrogel regarding particle size. Alhydrogel particles typically fall within the range of 1-10 microns^14^ whereas CoVaccine HT™ is typically showing droplet sizes around 130 nm^15,16^. Smaller particle sizes offer increased stability and enhanced adjuvanticity and in comparison, particle sizes of other commercial stable oil-in-water emulsion adjuvants (MF59 and AS03) are in the range of 160nm^17^. These oil-in-water emulsion adjuvants utilize squalene, a shark fat derived product ^9,18,19^. The use of squalane in CoVaccine HT™ as a plant-derived product may be advantageous due to availability, reduced regulatory burden, and potentially also ideologically to the population being immunised. In summary, CoVaccine HT™ could provide a distinct advantage over Alhydrogel as the more conventional adjuvant choice.

Here we tested the immunogenicity of SARS-CoV-2 Spike S1 proteins adjuvanted with either CoVaccine HT™, Alhydrogel, or phosphate buffered saline (PBS) in BALB/c mice. We assessed overall antibody titres, immunoglobulin subclass diversity, cell mediated immunity, and in-vitro neutralisation of wild-type SARS-CoV-2 virus. We demonstrate that CoVaccine HT™ elicits rapid humoral responses, increased subclass diversity, more interferon gamma (INFγ) production, and higher neutralising antibody titres than the other adjuvants. Collectively, CoVaccine HT™ may be advantageous over other adjuvants for a SARS-CoV-2 vaccine.

## 2. Methods

### 2.1 Vaccination and Serum Collection

BALB/c mice (7-8 weeks of age, male and female) were immunised twice, three weeks apart, intramuscularly (IM) with 5 μg of SARS-CoV-2 Spike S1 (Sino Biological 40592-V05H) protein with or without adjuvants, or adjuvant alone, using an insulin syringe with a 29-gauge needle. The adjuvants used were CoVaccine HT™ (Protherics Medicines Development Ltd, London, United Kingdom), or 2% Alhydrogel adjuvant (InvivoGen, San Diego, CA). Sera were collected by tail bleeding at 2 weeks post-immunisation or cardiac puncture for terminal bleeds. An additional serum sample was collected by cardiac puncture at day 28 along with splenocytes from three animals in the Spike S1 + CoVaccine HT™ (S1+CoVac) and S1 + Alum groups, and two animals in the S1+PBS group.

### 2.2 Serological Immunoglobulin Assays

Internally dyed, carboxylated, magnetic microspheres (Mag-PlexTM-C) were obtained from Luminex Corporation (Austin, TX, USA). The coupling of individually addressable microspheres with all previously mentioned proteins were conducted as described previously^20,21^. Microspheres dyed with spectrally different fluorophores were also coupled with bovine serum albumin as a negative control. SARS-CoV-2, SARS-CoV, and MERS-CoV specific immunoglobulin antibody titres in mouse sera were measured using a microsphere immunoassay as previously described with some minor alterations^13,22,23^. Briefly, microspheres coupled to his-tagged Spike S1 proteins of SARS-CoV-2, SARS-CoV, or MERS-CoV (Sino Biological 40591-V08H, 40150-V08B1, & 40069-V08H, respectively), and control beads coupled to bovine serum albumin (BSA) were combined and diluted in MIA buffer (PBS-1% BSA-0.02%Tween20) at a dilution of 1/200. Multiplex beads (at 50 μL containing approximately 1,250 beads of each type) were added to each well of black-sided 96-well plates. 50 μL of diluted serum were added to the microspheres in duplicate and incubated for 3 hours on a plate shaker set at 700 rpm in the dark at 37°C. The plates were then washed twice with 200 μL of MIA buffer using a magnetic plate separator (Millipore Corp., Billerica, MA). 50 μL of red-phycoerythrin (R-PE) conjugated F(ab’)2 fragment goat anti-mouse IgG specific to the Fc fragment (Jackson ImmunoResearch, Inc., West Grove, PA) were added at 1 μg/ml to the wells and incubated for 45 minutes. Antigen-specific IgG subclass titres were determined using mouse antisera at a 1:1000 dilution. Detection antibodies were subclass specific goat anti-mouse polyclonal R-PE-conjugated antibodies (Southern Biotech) used at a 1:200 dilution. The plates were washed twice, as described above, and microspheres were then resuspended in 120 μl of drive fluid (MilliporeSigma) and analysed on the MAGPIX Instrument (MilliporeSigma). Data acquisition detecting the median fluorescence intensity (MFI) was set to 50 beads per spectral region. Antigen-coupled beads were recognized and quantified based on their spectral signature and signal intensity, respectively. Assay cut-off values were calculated first by taking the mean of technical duplicate values using the average MFI (indicated as a dashed black line) from the adjuvant only control group. Cut-offs were generated by determining the mean MFI values plus three standard deviations as determined by Microsoft Office Excel program. Graphical representation of the data was done using Prism, Graphpad Software (San Diego, CA).

### 2.3 Plaque reduction neutralisation test (PRNT)

A PRNT was performed in a biosafety level 3 facility (at BIOQUAL, Inc.) using 24-well plates. Mouse sera pooled from individual mice within each group, were diluted to 1:10, and a 1:3 serial dilution series was performed 11 times. Diluted samples were then incubated with 30 plaque-forming units of wild-type SARS-CoV-2 (USA-WA1/2020, BEI Resources NR-52281) in an equal volume of culture media (DMEM-10% FBS with gentamicin) for 1hr at 37°C. The serum-virus mixtures were added to a monolayer of confluent Vero E6 cells and incubated for 1 hour at 37°C in 5% CO_2_. Each well was then overlaid with 1mL of culture media containing 0.5% methylcellulose and incubated for 3 days at 37°C in 5% CO_2_. The plates were then fixed with methanol at −20°C for 30 minutes and stained with 0.2% crystal violet for 30 minutes at room temperature. Neutralisation titres were defined as the highest serum dilution that resulted in 50% (PRNT_50_) and 90% (PRNT_90_) reduction in the number of plaques.

### 2.4 Preparation of mouse splenocytes and FluoroSpot assay

Mouse spleens were harvested at day 7 after the second dose, minced, passed through a cell strainer, and cryopreserved after lysis of red blood cells. Cellular immune responses were measured by IFN-γ FluoroSpot assay according to the manufacturer’s instructions (Cat. No. FSP-4246-2 Mabtech, Inc., Cincinnati, OH). Briefly, splenocytes were rested at 37°C, in 5 % CO_2_ for 3 hours after thawing to allow removal of cell debris. A total of 2.5 × 10^5^ cells per well in serum-free CTL-Test™ medium (Cellular Technology Limited, Shaker Heights, OH) were added to a 96 well PVDF membrane plate pre-coated with capture monoclonal antibodies and stimulated for 40 hours with peptides, PepTivator^®^ SARS-CoV-2 Prot_S1 peptide pool consisting of 15-mer peptides with 11 amino acids overlapping, covering the N-terminal S1 domain of the Spike protein of SARS-CoV-2 (Miltenyi Biotec, Auburn, CA) at 0.2 μg/mL and 0.5 μg/mL per peptide, or medium alone. Splenocytes at (5 × 10^4^ per well) were incubated with PMA (0.01 μM) /Ionomycin (0.167 μM) cocktail (BioLegend, San Diego, CA) as a positive control. The tests were set up in duplicates, and the costimulatory anti-CD28 antibody (0.1 μg/mL) was added to the cells during the incubation. Plates were developed using specific monoclonal detection antibodies and fluorophore-conjugated secondary reagents. Finally, plates were treated with a Fluorescence enhancer (Mabtech) to optimize detection and then air-dried. The spots were enumerated using the CTL ImmunoSpot® S6 Universal Analyzer (Cellular Technology Limited, CTL, Shaker Heights, OH), and the number of antigen specific cytokine-secreting spot forming cells (SFCs) per million cells for each stimulation condition was calculated by subtracting the number of spots detected in the medium only wells.

## 3 Results

### Murine immunisation with SARS-CoV-2 Spike S1 proteins

Neutralising antibodies of SARS-CoV-2 largely target the receptor binding domain present within the Spike S1 protein ^24^. Therefore, BALB/c mice were given two doses of commercially available Spike S1, 21 days apart (Fig.1A). To test whether adjuvants may alter immunological responses to the immunogen, mice were divided into four groups based on vaccine formulation. The mice receiving S1 protein and CoVaccine HT™ (S1+CoVac), Alhydrogel (S1+Alum), or PBS (S1+PBS) received SARS-CoV-2 Spike S1 mixed with either CoVaccine HT™ (“CoVac”), Alhydrogel (“Alum”), or PBS, respectively. One group received CoVaccine HT™ alone as an adjuvant control (Fig. 1A).

**Figure 1.**
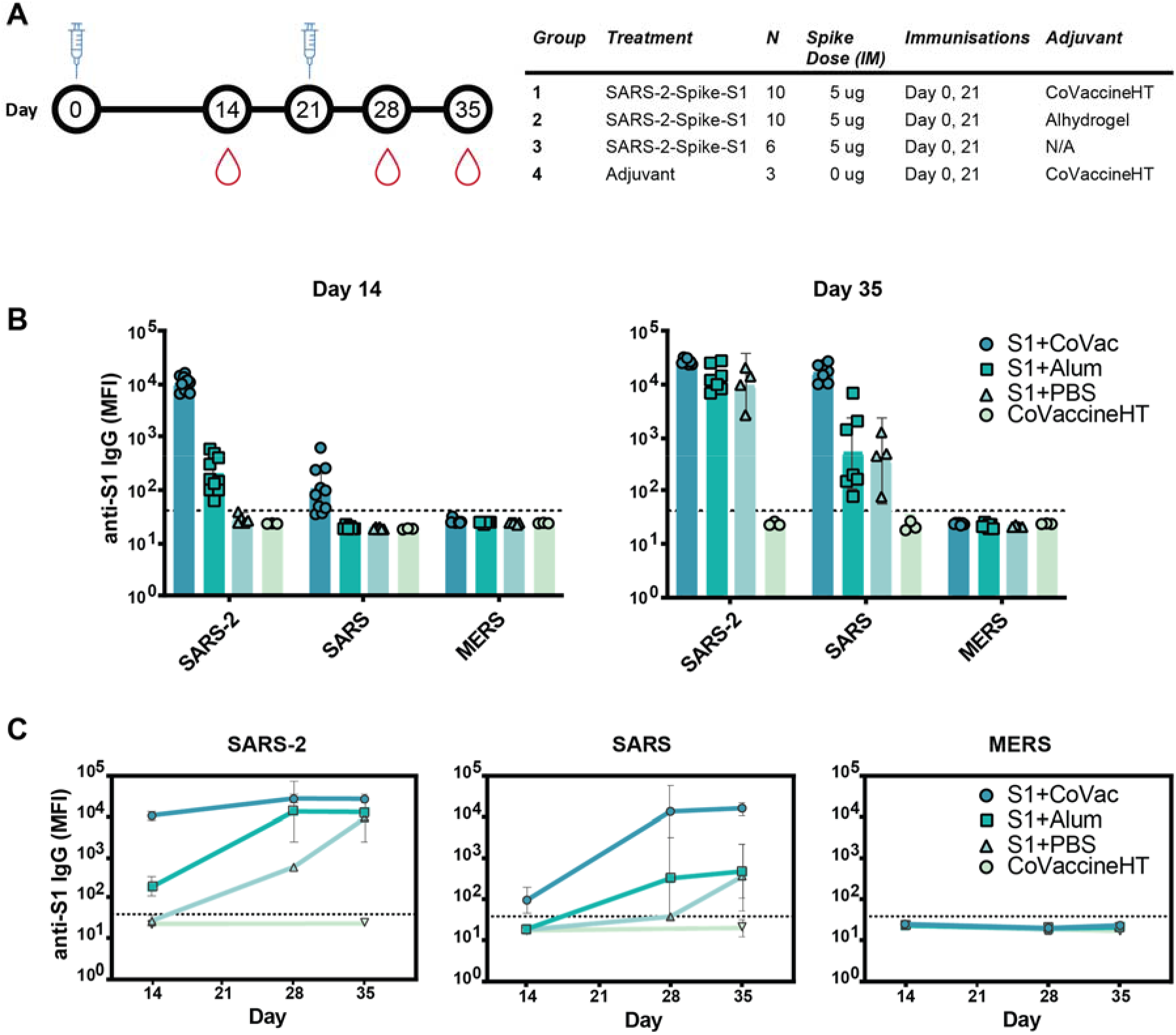
Immunogenicity and specificity to SARS-CoV-2 S1 immunisation. **A** Timeline schematic of BALB/c immunisations and bleeds with a table detailing the study design. **B** Median fluorescence intensity (MFI) of serum antibodies from each group binding to custom magnetic beads coupled with Spike S1 proteins from either SARS-CoV-2 (SARS-2), SARS-CoV (SARS), or MERS-CoV (MERS) on day 14 and 35. **C** Antibody reactivity to SARS-2, SARS, and MERS antigens throughout the study. Graphs in panels (B) and (C) are on a logarithmic scale representing geometric mean MFI responses with 95% confidence interval (CI). The dashed lines represent assay cut-off values determined by the mean plus three standard deviations of the negative control (BSA coupled beads).

### Adjuvants alter immunogenicity and specificity to immunisation

Serum analysis revealed high reactivity of SARS-CoV-2 S1 specific IgG antibodies in S1+CoVac after a single dose while S1+Alum titres were near baseline (Fig.1B). Only one animal showed a detectable titre in the antigen alone group at this time point. Only in the group with CoVaccine HT™ a low level of cross reactivity was observed after the first dose to SARS-CoV S1. On day 35, S1+Alum and S1+PBS displayed significantly higher antibody responses compared to day 14 and variations among individual animals were reduced. S1+CoVac treated animals on day 35 consistently showed very high antibody responses in every animal. Similarly, cross-reactivity with SARS-CoV S1 was greatly increased for all groups on day 35 (Fig.1B). As expected, due to its much lower sequence homology, the SARS-CoV-2 S1 did not induce IgG responses to MERS-CoV S1.

In patients suffering from COVID-19, high RBD-specific IgG titres have been observed^25^. However, higher titres of SARS-CoV-2 Spike-specific IgG are associated with patients that did not require intensive care unit treatment while lower titres are associated with increased disease severity^26^. Therefore, the antibody response kinetics may be an important factor for a successful vaccine candidate. Time-course analysis of IgG responses reveal that adjuvanted S1 may be crucial for strong, early IgG responses with SARS-CoV-2 specificity while a second dose may decrease variability among individual animals and increase cross-reactivity (Fig.1C).

### CoVaccine HT™ improves IgG titres to SARS-CoV-2 and SARS-CoV S1 proteins

To further investigate the matured IgG responses, sera from day 35 were titrated in a four-fold dilution series starting at 1/250 and analysed by microsphere immunoassay (MIA). The S1+Alum and S1+PBS groups showed reactivity to SARS-CoV-2 S1 when diluted up to 1/256,000, indicating an abundance of antigen-specific IgG in the sera (Fig. 2A). Titrating sera from S1+CoVac however, revealed saturating levels of IgG for five dilutions and detectable IgG levels were present down to a 1/65.5 million dilution. Antiserum to S1+CoVac also showed significantly greater cross reactivity to SARS-CoV S1 compared to the other groups. All groups remained negative for cross reactivity to MERS-CoV S1 (Fig.2A). These data suggest that immunisation with SARS-CoV-2 S1 and CoVaccine HT™ elicits robust antigen-specific IgG response with the expected cross-reactivity profile to include SARS-CoV S1.

**Figure 2.**
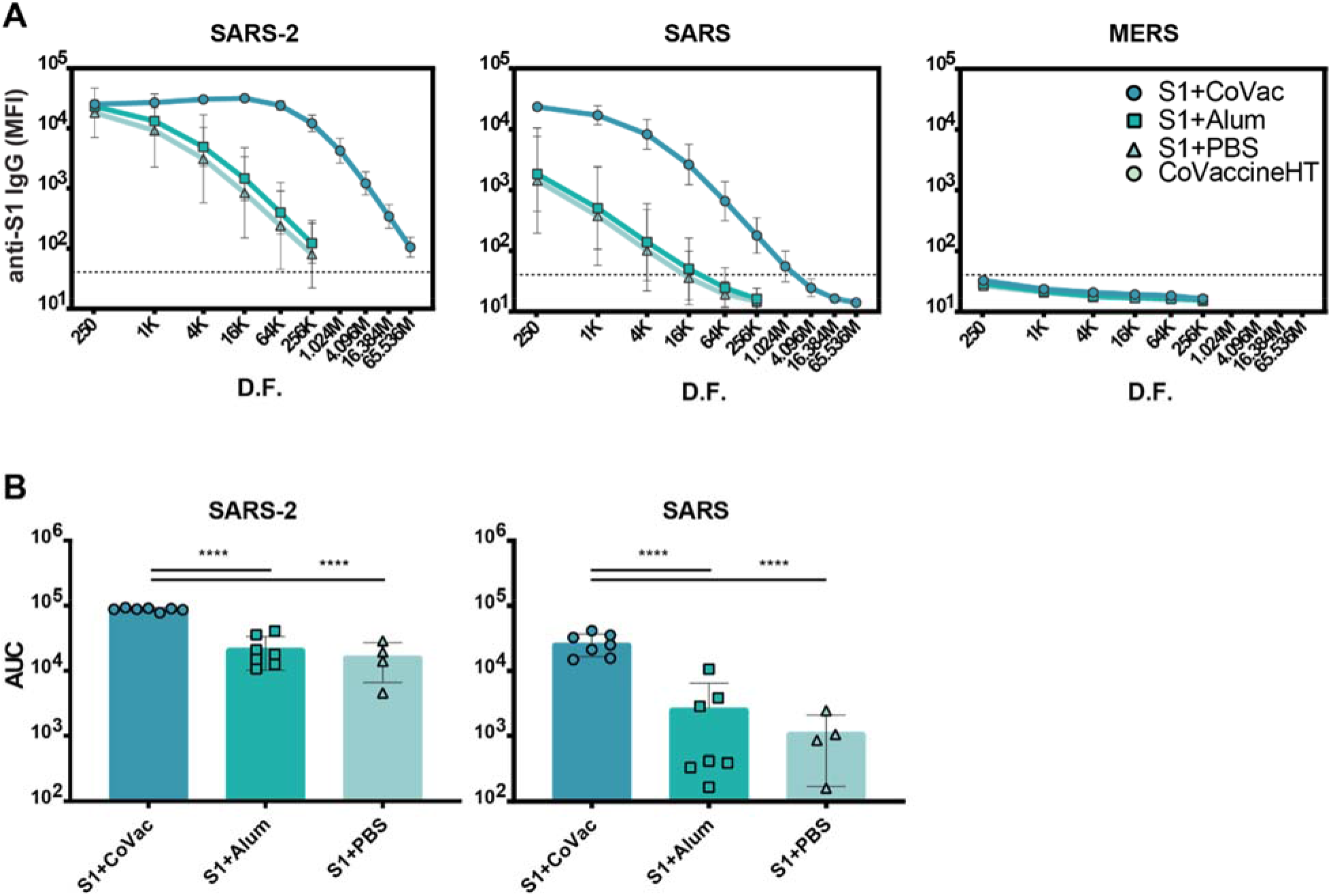
Serum IgG titres against Coronavirus S1 antigens. **A** Antigen reactivity in a four-fold dilution series of mouse sera. **B** Area under the curve (AUC) of data in (A). Both graphs are in log scale with geometric mean and 95% CI. The dashed lines in panel (A) represent the cut-off value determined by the mean plus three standard deviations of the negative control (BSA coupled beads). K= x1000, M= x1,000,000, D.F.= dilution factor. Statistics by standard one way-ANOVA. **** = p-value < 0.0001.

### Increased IgG subclass diversity and enhanced viral neutralising antibody titres with CoVaccine HT™

Adjuvants serving as TLR4 agonists, such as postulated for CoVaccine HT™, elicit a primarily T_h_1 type response ^27,28^. Meanwhile, Alhydrogel facilitates a mainly T_h_2 type response, possibly through NOD-like receptor signalling ^29,30^. IgG subclass analysis can be used to determine if a T_h_1 or T_h_2 response may have been more prominent. Therefore, sera from each S1+adjuvant group were analysed for their subclass composition (Figure 3). Consistent with previous findings, the S1+CoVac group displayed a diverse immunoglobulin response composed of IgG1, IgG2a, and IgG2b subclasses all of which were further elevated after a second dose of vaccine. Low levels of IgG3 were also observed. Alternatively, the Alum and antigen alone groups primarily produced an IgG1 response with some detectable IgM in the Alum group, representing a classical T_h_2-biased humoral response. Heterogeneous subclass populations such as those observed in the S1+CoVac group are typically associated with T_h_1 responses while IgG1 is characteristic of a T_h_2 response. To further investigate the nature of these adjuvant effects, the subclass data were stratified to analyse ratios of T_h_1 vs T_h_2 subclasses (Figure 3C). This analysis clearly shows that of the three tested formulations, only S1+CoVac induced a relatively balanced humoral response. Furthermore, only the S1+CoVac formulation was able to induce detectable SARS-CoV-2 neutralising antibody titres as demonstrated in a plaque reduction neutralisation test using wildtype virus (Table 1). PRNT_90_ and PRNT_50_ titres for this formulation indicate potent neutralisation (1:1620).

**Table 1.** SARS-CoV-2 neutralisation titres.

| Group ID | Titre (PRNT<sub>90</sub>) | Titre (PRNT<sub>50</sub>) |
| --- | --- | --- |
| S1+CoVac | 1620 | 1620 |
| S1+Alum | <20 | <20 |
| S1+PBS | <20 | <20 |
| CoVaccine HT | <20 | <20 |

**Figure 3.**
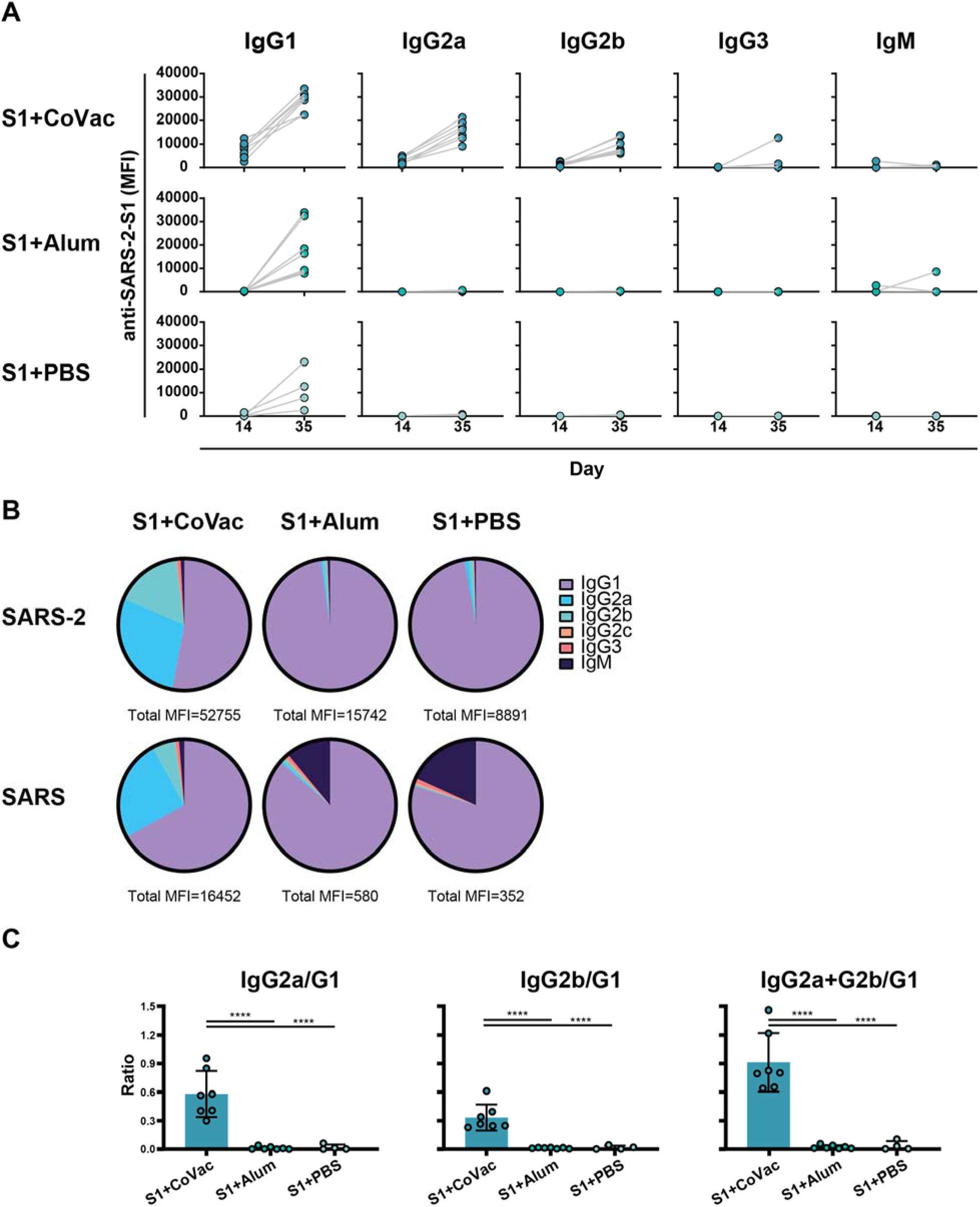
Adjuvant effects on immunoglobulin subclass diversity. **A** IgG subclasses reacting with SARS-2 S1 antigen between day 14 and day 35 plotted on a linear scale. **B** Relative abundance of Immunoglobulin isotypes and IgG subclasses reacting to SARS-2 and SARS antigens determined by subtracting the specified subclass cut-off values from the geometric mean of each group. The total MFI from which the subclasses are a fraction of is listed below each pie-chart. **C** Ratios of subclasses. The normalized MFI values of each subclass per mouse were plotted as ratios using mean and SD. Statistics by standard one way-ANOVA. **** = p-value < 0.0001.

### Adjuvant effect on the SARS-CoV-2 S1-specific INFγ responses

We assessed the adjuvant effect of CoVaccine HT™ and Alum on the cellular immune responses directed against SARS-CoV-2 S1 using an IFN-γ FluoroSpot assay. Individual mouse spleens from each group harvested at day 7 post-second immunisation were processed, and single cell suspensions stimulated with SARS-CoV-2 S1 peptides. The number of IFN-γ secreting cells from the mice given CoVaccine HT™ was significantly higher than those for mice given Alum or S1 antigen only at two different peptide concentrations (Figure 4). Splenocytes from unvaccinated (naïve) mice did not respond to S1 peptide stimulation with only 2 spot forming cells (SFCs)/10^6^ cells detected. The results suggest that CoVaccine HT™ is a superior adjuvant for induction of an antigen-specific Th1-focused cellular immune response, which is critical for SARS-CoV-2 vaccine development.

**Figure 4.**
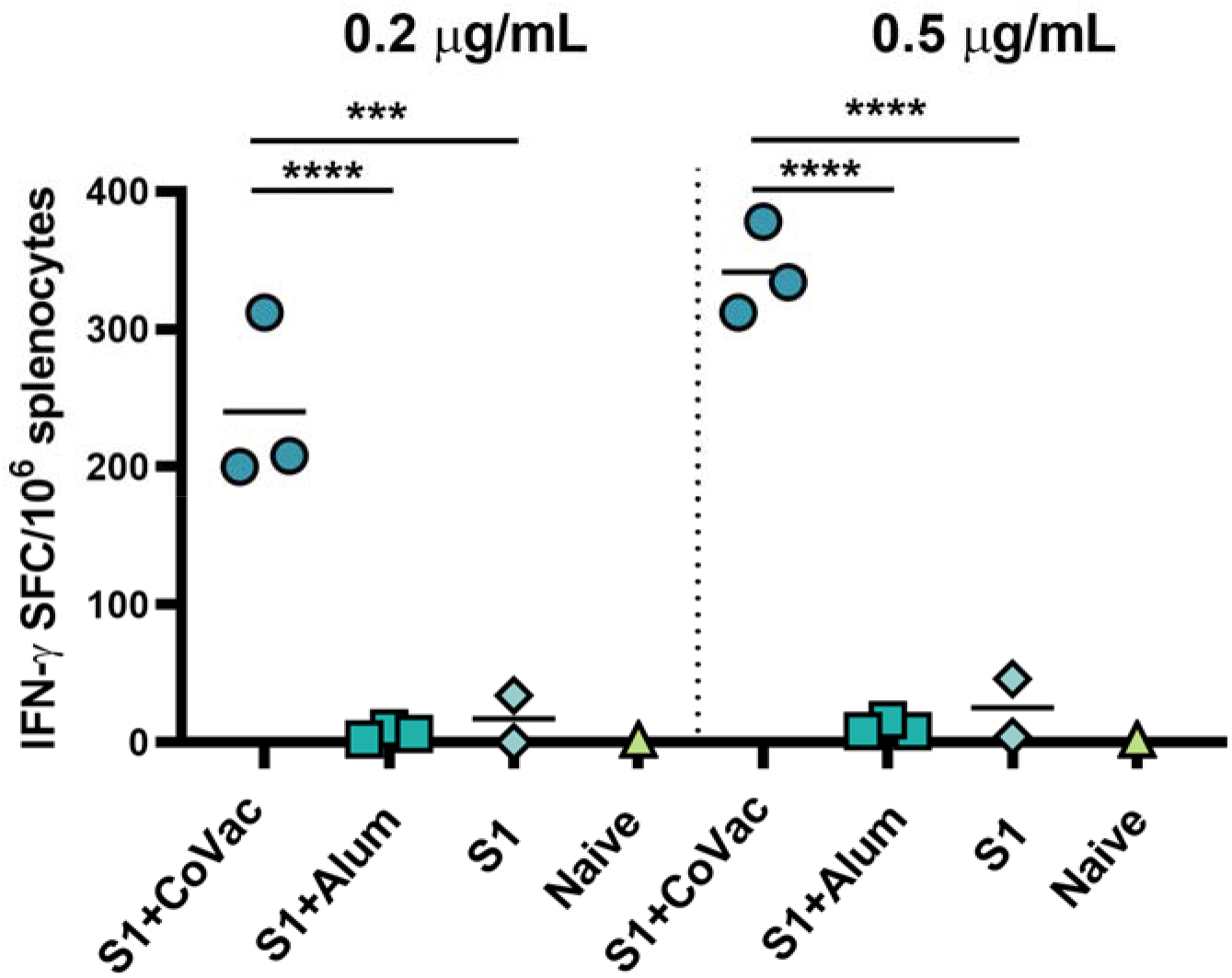
Detection of IFN-γ secreting cells from mice immunised with SARS-CoV-2 vaccines. The splenocytes were obtained from mice (2 to 3 per group) immunised with SARS-CoV-2 S1 protein, adjuvanted with CoVaccine HT™ or Alum, or S1 protein alone on day 28 (one-week after booster immunisations). Pooled splenocytes obtained from two naïve mice were used as controls. The cells were incubated for 40 hours with PepTivator® SARS-CoV-2 Prot_S1 peptide pools at 0.2 μg/mL or 0.5 μg/mL per peptide or medium. IFN-γ secreting cells were enumerated by FluoroSpot as detailed in the methods section. The results are expressed as the number of spot forming cells (SFC)/106 splenocytes after subtraction of the number of spots formed by cells in medium only wells to correct for background activity. *** p ≤ 0.001, **** p ≤ 0.0001.

## 4. Discussion

The COVID-19 pandemic has stimulated global efforts to rapidly develop vaccines against SARS-CoV-2. Many vaccine strategies are being explored, including inactivated virus, non-replicating viral vectors, recombinant protein, DNA, and RNA several of which have reached human clinical trials ^7,6^. The number of clinically applied adjuvants are limited and include Alum and newer formulations such as MF59 and AS03, both oil-in-water emulsions using squalene ^31^. The small number of adjuvants approved for clinical use has limited vaccine development in the past and impacts current clinical trials of SARS-CoV-2 vaccines. Many approaches use no adjuvant, Alum, MF59, or AS03, however, Novavax is testing the experimental adjuvant Matrix-M™ ^32^. While Alum is known to primarily enhance a T_h_2 response, Matrix-M™ and CoVaccine HT™ have both been shown to elicit a T_h_1 response with recombinant subunits. Due to the previously observed potential for enhanced immunopathology associated with primarily T_h_2-targeted anti-SARS-CoV vaccines^33,34^, the development of a COVID-19 vaccine may require testing of a multitude of adjuvants to elicit protective immune responses to SARS-CoV-2. The squalane-in-water based adjuvant, CoVaccine HT™, has previously been shown to induce potent virus neutralisation antibody titres and protective efficacy in mice and non-human primates to several infectious agents, and has recently been licensed by Soligenix, Inc. for use in SARS-CoV-2 vaccine development ^35–38^.

In this study, we investigated the immunogenicity of recombinant SARS-CoV-2 Spike S1 alone or in combination with Alum or CoVaccine HT™ as potential adjuvants. Overall, we observed the most potent humoral and cellular immune responses, including neutralising antibody responses in the CoVaccine HT™ study group. Day 14 titres (post-dose 1) indicate that this formulation may even be efficacious after administration of a single dose, however, this was not investigated in the current study. Neither antigen alone nor the combination with Alum was able to induce detectable neutralising antibodies with the model antigen utilized in this study. This may be due to slower response kinetics caused by antigen presentation, subclass homogeneity, or T_h_2 restricted immune responses compared to administering S1 with CoVaccine HT™.

Immunogenicity of protein subunit vaccines is often inferior in generating robust immune responses compared to other platforms such as those based on live attenuated viruses. As seen here, the (monomeric) S1 domain alone is not adequate for generating a high titre immune response. The addition of CoVaccine HT™ improved antibody titres and response kinetics and proved to induce high titres of antibodies neutralising wild-type SARS-CoV-2. It has been shown by others that SARS-CoV-2 S1 IgG titres correlate with viral neutralisation in humans^39^. Virus neutralising responses in rabbits after two immunisations with 50μg of SARS-CoV-2 S1 and Emulsigen adjuvant (oil-in-water emulsion) were at 1:160 in the wild-type neutralisation assay, compared to titres at 1:800 achieved when immunising with SARS-CoV-2 RBD ^40^. Similarly, emulsion-based adjuvants improved kinetics and antibody titres in guinea pigs compared to Alhydrogel or no adjuvant with HIV-1 gp140 immunisation^41^. This demonstrates the importance of achieving an antigen/adjuvant combination with desirable properties. In our study, post-dose 1 titres in the S1+CoVac group resemble post-dose 2 titres with Alum or no adjuvant and may suggest at least partial protection after a single dose. Generating potent immunity after a single dose is an attractive target for any SARS-CoV-2 vaccine in development and may improve the impact of a vaccine on the further course of the pandemic.

The high potency for SARS-CoV-2 S1 in the CoVaccine HT™ formulation may be attributable to the observed immunoglobulin subclass diversity. This indicates CoVaccine HT™ may efficiently induce class switching often considered to increase antibody affinity. Furthermore, a broad IgG subclass composition is key for inducing complement-mediated antibody effector functions as well as neutralisation and opsonisation, which are typically essential for mitigating viral infections. The ideal antibody population has yet to be elucidated for combating SARS-CoV-2. However, our murine serological data suggests kinetics and subclass diversity may be key to developing effective immune responses. Additionally, we have demonstrated that CoVaccine HT™ is not only a suitable adjuvant for vaccination but is preferable to Alhydrogel given the quality of the humoral response due to rapid onset, balance, overall magnitude of the response, as well as significantly greater cell-mediated immune responses.

Concerns have been raised regarding antibody dependent enhancement (ADE) with SARS-CoV-2 infection or immunisation. This phenomenon occurs when non-neutralising or poorly binding antibodies interact with Fc receptors on antigen presenting cells and facilitate infection. This interaction increases pro-inflammatory cytokine production which exacerbates immunopathology ^42^. ADE was previously observed with SARS-CoV infection caused by anti-Spike antibodies through the FcγR and FcγRII pathways ^43,44^. In respiratory syncytial virus infections, a T_h_2 response alone can lead to aberrant immune responses associated with ADE caused by either a prior infection or immunisation ^45,46^. For these reasons, caution may be warranted when using Alum as an adjuvant for SARS-CoV-2, at least for the subunit protein used in this study. Additionally, our data suggest that antigen alone may also drive a predominantly T_h_2 type response with recombinant S1 antigen immunisation in mice. In contrast, the subunit protein with CoVaccine HT™ produces neutralising antibodies while boosting T_h_1 responses which may increase durability, vaccine safety, and efficacy.

Previous studies of SARS-CoV, the most closely related human betacoronavirus to SARS-CoV-2, have shown that recovered patients develop substantial T cell responses, which persist for up to 11 years ^47,48^. In addition, animal studies indicated that T cell responses play a crucial role in protection against SARS-CoV infection^49–51^, suggesting that it is most likely that T cell responses to SARS-CoV-2 are protective. Despite very limited understanding on the adaptive immune responses to SARS-CoV-2, it has been reported that virus-specific T cell responses are detected in 70% to 100 % of COVID-19 convalescent patients and about 50% of the CD4+ T cell response is directed against S protein and correlates with the magnitude of anti-S antibody response^52^. Furthermore, the majority of CD4^+^ T cells appears to be T_h_1 type with little or no T_h_2 cytokine secretion detected^52^. This suggests that a SARS-CoV-2 vaccine candidate consisting of S protein can induce a robust CD4^+^ T cell response that recapitulates the elicited immune response during the natural infection. In this study we showed that SARS-CoV-2 S1 adjuvanted with CoVaccine HT™ elicits a strong IFN-γ secreting cellular immune response upon peptide stimulation, indicating the induction of a T_h_1-targeted T cell response, which highlights the potential of this antigen/adjuvant combination to protect against SARS-CoV-2 infection. However, further in-depth analysis of cellular immune responses will be needed to characterize CD4^+^ and CD8^+^ T cell responses and their correlation with the antibody titres.

Altogether, CoVaccine HT™ is an effective adjuvant that in combination with a properly chosen recombinant subunit protein promotes rapid induction of balanced humoral and cellular immune responses and allows accelerated preclinical and clinical development of a SARS-CoV-2 vaccine to mitigate the ongoing COVID-19 pandemic.

## Acknowledgements

The authors would like to acknowledge the following reagent deposited by the Centers for Disease Control and Prevention and obtained through BEI Resources, NIAID, NIH: SARS-Related Coronavirus 2, Isolate USA-WA1/2020, NR-52281. We further would like to acknowledge the gift of CoVaccine HT™ from Protherics Medicines Development (London, UK) and would like to thank Dr. Oreola Donini (Soligenix, Inc., Princeton, NJ) for an expanded collaboration including strategic discussions and critical reading of this manuscript. We would further like to acknowledge partial funding for these studies from R01AI132323 (National Institute of Allergy and Infectious Diseases), from P30GM114737 (Centers of Biomedical Research Excellence, National Institute of General Medical Sciences) and institutional funds.

## Author Contributions

Authors BKH and CYL have equal contribution.

BKH, CAW, TW, CYL, MML, AL: Conceived and designed the experiments. BKH, TW: Immunisations and blood collection. BKH, CYL: Splenectomy. BKH: Microsphere immunoassays and Flow cytometry studies. CYL: Fluorospot assay. BIOQUAL: Virus neutralisation assays. BKH, CAW, TW, MML, AL: Manuscript writing and editing.

